# Integrated transcriptional analysis of the cellular and extracellular vesicle RNA content of Candida auris in response to caspofungin

**DOI:** 10.1101/2020.12.04.411843

**Authors:** Daniel Zamith-Miranda, Rafaela F. Amatuzzi, Sharon T. Martins, Alexandre Z. Vieira, Isadora M. da Rocha, Marcio L. Rodrigues, Gabriel Trentin, Fausto Almeida, Ernesto S. Nakayasu, Joshua D. Nosanchuk, Lysangela R. Alves

## Abstract

*Candida auris* has emerged as a serious worldwide threat by causing invasive infections in humans that are frequently resistant to one or more conventional antifungal medications, resulting in high mortality rates. Against this backdrop, health warnings around the world have focused efforts on understanding *C. auris* fungal biology and effective treatment approaches to combat this fungus. To date, there is little information about *C. auris* gene expression regulation in response to antifungal treatment. Our integrated analyses focused on the comparative transcriptomics of *C. auris* in the presence and absence of caspofungin as well as a detailed analysis of the yeast’s extracellular vesicle (EV)-RNA composition. The results showed that genes coding oxidative stress response, ribosomal proteins, cell wall, and cell cycle were significantly upregulated in the presence of caspofungin, whereas transcriptional regulators and proteins related to the nucleus were downregulated. The mRNAs in the EVs were associated with stress responses induced by caspofungin and the ncRNA content of the EVs shifted during caspofungin treatment. Altogether, the results provide further insights into the fungal response to caspofungin and demonstrate that analyses of *C. auris* growth under antifungal stress can elucidate resistance and survival mechanisms of this fungus in response to medical therapy.

## INTRODUCTION

Invasive fungal infections are responsible for over 1.5 million deaths per year (1). Among these diverse fungal diseases, bloodstream infections caused by *Candida* species are the most common cause of invasive disease (2). Although *C. albicans* is frequently the most common species associated with invasive candidiasis (3), infections by non-albicans species have increased. These non-albicans species more frequently display multidrug resistance, representing a major problem in healthcare globally, particularly in immunocompromised hospitalized patients (4, 5).

The emerging multidrug resistant fungal pathogen *C. auris* is responsible for numerous nosocomial outbreaks in healthcare settings, from nursing homes to intensive care units, on all inhabited continents (6, 7). *C. auris* was first reported in 2009 as the etiological agent of an ear infection in Japan (8, 9), and it remains understudied. The difficulty in identifying the fungus by classic phenotyping and our gaps in understanding its propagation mechanisms in healthcare settings worldwide has impeded our capacity to control its spread (10). Due to these issues as well as its remarkable drug resistance, *C. auris* is the first fungal pathogen to be classified as a global public health threat (8). This fungus can be transmitted rapidly among patients, persist on hospital surfaces and medical devices (11, 12), and has a high mortality, particularly among patients who have undergone multiple medical interventions (10, 13–15).

*C. auris* has an extremely high frequency of multidrug resistance to polyenes, echinocandins, and azoles, especially fluconazole (16). Resistant phenotypes are currently being deciphered, although it is clear that polymorphisms in ERG11 and FKS1 genes are associated with the development of fluconazole and echinocandin resistance, respectively (16, 17). Another feature that may contribute to the resistance of *C. auris* to a variety of antimicrobial agents is biofilm formation (18). Even though it is not as thick or adherent to medical devices substrates compared to *C. albicans* biofilm (18), *C auris* biofilm confers drug resistance to fluconazole (18, 19) and caspofungin (18). This factor may be associated with an increased activity in efflux pumps (19). *C. auris* biofilm transcriptome analysis has revealed that genes coding efflux pumps such as ATP-binding cassette (ABC) and major facilitator superfamily (MFS) transporters are upregulated in the presence of antifungal agents (19).

Despite the increased *C. auris* medical importance of *C. auris*, little is known about its mechanisms of pathogenicity and virulence (4). The *C. auris* genome has many potential virulence factors such as proteinases, lipases, phospholipases, adhesins, efflux pumps, biofilm formation and drug transport, involved with azole resistance (20). Pathogenic fungi secrete a myriad of immunoreactive molecules, cytotoxic proteins, virulence factors and a subset of functional RNAs to assist them in invasion and survival in the host environment (21, 22). Recently, studies have revealed that extracellular vesicles (EVs) can deliver these functional molecules to target cells (23, 24). EVs are lipid bilayer structures released by organisms from all domains of life (25, 26). EVs from several pathogenic fungi species play diverse biological roles related to cell-cell communication and pathogen-host interaction (27–31). To date, there is evidence that suggests an association between fungal EVs and antifungal response, however, more studies need to be performed in order to clarify this matter (32).

The echinocandins are the drugs of choice for the initial treatment of *C. auris*, as recommended by the CDC (https://www.cdc.gov/fungal/candida-auris/c-auris-treatment.html), so in this study we characterized the alteration in RNA abundances of yeasts treated with the antifungal caspofungin and correlated these findings with the RNA content of EVs derived from the same cells. Our data showed that caspofungin not only affected cell morphology, but also led to a transcriptional response with upregulation of cell wall associated processes, and silencing of pathways that are associated with cellular responses to stress. Our study contributes with new information on *C. auris* that highlights genetic responses to a standard antifungal and how the cell can circumvent the treatment to survive and persist in the host.

## MATERIAL AND METHODS

### Fungal growth conditions

*C. auris* strains MMC1 and B8441 (South Asia Clade) were maintained at −80°C. B8441 was obtained from the CDC (also known as CDC387). After thawing in Sabouraud broth, suspensions were incubated at 30□C for 24 hours. Yeast cell suspensions were then plated onto Sabouraud agar plates and incubated at 30 C for 48 hours. The plates were then stored at 4□C (for no longer than 4 weeks) and used in experiments. Yeast cell viability prior to experimental use and during culturing was monitored by propidium iodide (PI) staining and analysis by flow cytometry.

Based on MICs, growth curves of *C. auris* in the presence of caspofungin were performed with concentrations of 12.5 ng/mL and 100 ng/mL for lines MMC1 and B8441, respectively. The assays were performed in technical and biological triplicates in a 96-well flat bottom translucent plate to a final volume of 200μL per well and 5×10^5^ cells/mL. The cells were incubated in a microplate reader (Synergy Biotek) for 72h at 30°C, with optical density (OD) readings taken every 1h with a wavelength of 540nm, with prior shaking of the plate for 30 seconds before each reading. A non-linear model, Gompertz curve, was used to analyze the growth curves, the significance was calculated by ANOVA, comparing control versus treatment. The results represent the mean ± standard deviation of three independent experiments. The curves and analyzes were performed using the GraphPad Prism 8.0 program.

For Western blot analysis, protein extracts were separated by SDS-PAGE in a 15% polyacrylamide gel and transferred to a nitrocellulose membrane. The membrane was blocked and incubated with the primary antibodies mouse anti-S6 1:700 dilution, rabbit anti-H4 1:800 dilution. The secondary antibodies used were goat IRDye680-conjugated anti-rabbit or mouse IgG (Sigma) diluted 1:10,000. The result was visualized in an Odissey Imaging System (Li-Cor). The intensity of the bands was calculated using ImageJ software.

### Scanning electron microscopy (SEM)

The cells were collected by centrifugation at 5,000 rpm for three minutes and were washed 3 times with PBS buffer and then was fixed with 1 mL of 2.5% gluteraldehyde in 0.1M cacodylate buffer, pH 7.2 for 1 hour at room temperature and subsequently they were washed 3 times with post-fixation solution (0.1M cacodylate buffer, 0.2M sucrose and 2mM MgCl_2_). After the fixation step, the cells were adhered in round coverslips with poly-I-lysine for 30 minutes. The excess was quickly removed, and the samples were gradually dehydrated, by adding sequentially adding ethanol 30%, ethanol 50%, ethanol 70% for 5 minutes each and 95% ethanol, 100% ethanol, twice, for 10 minutes each. After dehydration, 100% ethanol was replaced with carbon dioxide (CO_2_) with a 10 times cycle. At the end of the process, the coverslips were metallized with gold and visualized in a scanning electron microscope (Jeol JSM-6010 Plus-LA) at 5 Kv.

### Cell wall staining

Yeast cells, in Sabouraud broth, were treated or not with caspofungin at a concentration of 12.5ng/mL (MMC1 – MIC = 2 μg/mL) and 100ng/mL (B8441 – MIC = 250 ng/mL) for 24 hours at 37□C. Cells were washed with PBS, fixed with paraformaldehyde 4% for 30 minutes at room temperature (RT). After washing again, cells were blocked with BSA 1% in PBS for 1 hour at RT and then incubated with concanavalin A (5 μg/mL), wheat-germ agglutinin (WGA – 10 μg/mL) and uvitex 1% for 30 minutes at RT. After washing with PBS, samples were analyzed under an Observer Z1 (Zeiss) microscope. In parallel, cells that were only stained with concanavalin A were quantified in a flow cytometer (FacScalibur) for quantification.

### Production of reactive oxygen species

Yeast cells (5 x 10^6^ per well) were incubated with 50 μM of H2DCFDA for 20 minutes at RT. Cells were washed with PBS and incubated or not with caspofungin in DMEM/F12 for 3 hours at 37 □C. Cells were washed with PBS and analyzed by flow cytometry (FacScalibur).

### Extracellular vesicles isolation

One colony from each strain was inoculated into Sabouraud broth for 24 hours at 37°C at 200 rpm. Cell density was adjusted to 10^6^ cells/mL in a total volume of 400 mL with or without the addition of caspofungin at a concentration of 12.5ng/mL (MMC1 – MIC = 2 μg/mL) and 100ng/mL (B8441 – MIC = 250 ng/mL). After 24 hours at 37°C at 200 rpm, the cells were centrifuged for two cycles, first at 8000 rpm for 15 minutes at 4°C and the second at 14000 rpm for 15 minutes at 4°C. Then, the supernatant was filtered on a 0.45 μm membrane (Milipore) followed by concentration of the supernatant using an Amicon with a 100KDa cutoff membrane. The concentrate was centrifuged two times for 1 hour at 150,000g at 4°C with a washing step in between. The supernatant was discarded and the EVs were suspended in 100μL of PBS.

### Nanoparticle tracking analysis (NTA)

Nanoparticle Tracking Analysis (NTA; Nanosight LM-10 −Malvern Panalytical) was used to determine the concentration and size of EVs isolated from the *C. auris* strains samples. EV samples were diluted in PBS prior to injection. The videos were set to three runs each of 60 s, and the detection threshold was defined as 2 and camera level as 9. The data were analyzed using NTA 3.1 software. Total yield (EV particles/mL) was calculated based on dilution factors. Statistical analysis was performed with Minitab Statistical Software 17.0, where samples were submitted to One-way variance analysis, and values were compared with Tukey test with 5% probability.

### Transmission electron microscopy

A total of 50 μl of EVs samples were deposited onto carbon-coated 300-mesh copper grids, incubated for 60 min at room temperature, washed with PBS, fixed with Karnovisk solution for 10 min, and then washed 3 times with cacodylate buffer 0.1M. The grids were then stained with 30 μl of 5% (v/v) uranyl acetate. Excess solution was blotted off and the grids were washed with ultrapure water 18.2Ω and dried overnight. Images of the EV were captured using a JEOL JEM 1400 electron microscope (JEOL Ltd) operated at 100 kV and magnification at 20k or 25k.

### RNA isolation and sequencing

Total RNA was isolated from 1 x 10^7^ *C. auris* cells using the miRCURY RNA isolation kit (Exiqon - Qiagen) with adaptations. A 1:1 volume of glass beads was added to the lysis buffer along with the yeast cells, and the mixture was subjected to 10 rounds of 1 min at 4□C of vortex agitation in order to disrupt the fungal cell wall. After centrifugation, total RNA was isolated according to the manufacturer’s instructions.

The small RNAs from the yeast cells (1 x 10^7^) were isolated with the miRNeasy kit (Qiagen) with modifications. A 1:1 volume of glass beads was added to the trizol buffer, followed by 10 rounds of 1 min at 4□C of vortex agitation in order to disrupt the cell wall. After centrifugation, small RNAs were collected according to the manufacturer’s instructions.

EV RNA isolation was carried out using the miRNeasy kit (Qiagen) according to the manufacturer’s instructions. The DNA cleanup step was performed with all samples using the RNAse-free DNAse protocol (Qiagen). For RNA quantification and integrity analysis we used a Qubit fluoremeter (Thermo Fisher) and an Agilent 2100 Bioanalyzer; RNA 6000 pico and RNA small kits (Agilent Technologies).

For the cellular RNA, the sequencing library was constructed with the TruSeq Stranded mRNA kit (Illumina) prepared according to the manufacturer’s instructions and all the samples were prepared in three independent replicates. For the small RNA and EV RNA, the libraries were constructed using the TruSeq small RNA kit (Illumina). The libraries were prepared according to the manufacturer’s instructions with slight adaptations. In the step of size-selecting the samples by using the acrylamide gel, instead of cutting between custom markers, we cut the molecular sizes corresponding to mRNAs as well as smaller ones. The samples were prepared in three independent replicates and RNAseq was performed on a HiSeq 2500 (Illumina, single-end 50-bp SR mid output run) at the Life Sciences Core Facility (LaCTAD), a part of the University of Campinas (UNICAMP).

### qRT-PCR

For the quantitative real time PCR the experimental design was performed according to the Minimum Information for Publication of Quantitative Real-Time PCR Experiments (MIQE) guidelines (33). Total RNA was isolated in duplicate from 1 x 10^7^ *C. auris* cells using the miRCURY RNA isolation kit (Exiqon - Qiagen) with adaptations. A 1:1 volume of glass beads was added to the lysis buffer along with the yeast cells, and the mixture was subjected to 10 rounds of 1 min at 4□C of vortex agitation in order to disrupt the fungal cell wall. After centrifugation, total RNA was isolated according to the manufacturer’s instructions and quantified using Qubit™ fluorometer RNA HS kit (Thermo Fisher) and the RNA integrity was assessed with Bioanalyzer RNA PICO 6000 (Agilent). After isolation, 1 ug (for cellular RNA) and 20 ng (for EV RNA) was treated with 1 U of DNAse I RNAse-free (#EN0521 PROMEGA) according to manufacturer instructions. After that, the cDNA was synthesized from 1 μg of cellular RNA or 20 ng of vesicular RNA as templates. For the reverse transcriptase reactions 0.3 μM random primer (Invitrogen) and 1 μl of reverse transcriptase (Superscript II, Thermo Scientific), according to the manufacturers’ instructions. PCR was performed with 40 ng of cDNA for the cell and 1.6 ng of cDNA as the template and GoTaq™ master mix according to manufacturer instructions (Promega). The oligonucleotides were designed with PRIMER-Blast using the following parameters: PCR product size maximum of 250 nt. Tm varying from 57 to 63 °C, RefSeq mRNA as a database and Candida auris as the organism. The primer sets used for PCR are described below. The qPCR was performed in four technical replicates for each sample. The following program was used in the Lightcycler 480 (Roche) equipment: initial denaturation at 95 °C for 15 min and 45 cycles of 95 °C for 15 s, 62 or 64 °C for 20 s and 72 °C for 45 s. The reference gene used was histone H4 and the target genes and the primers used are listed in Table 1.

**Table 1.**
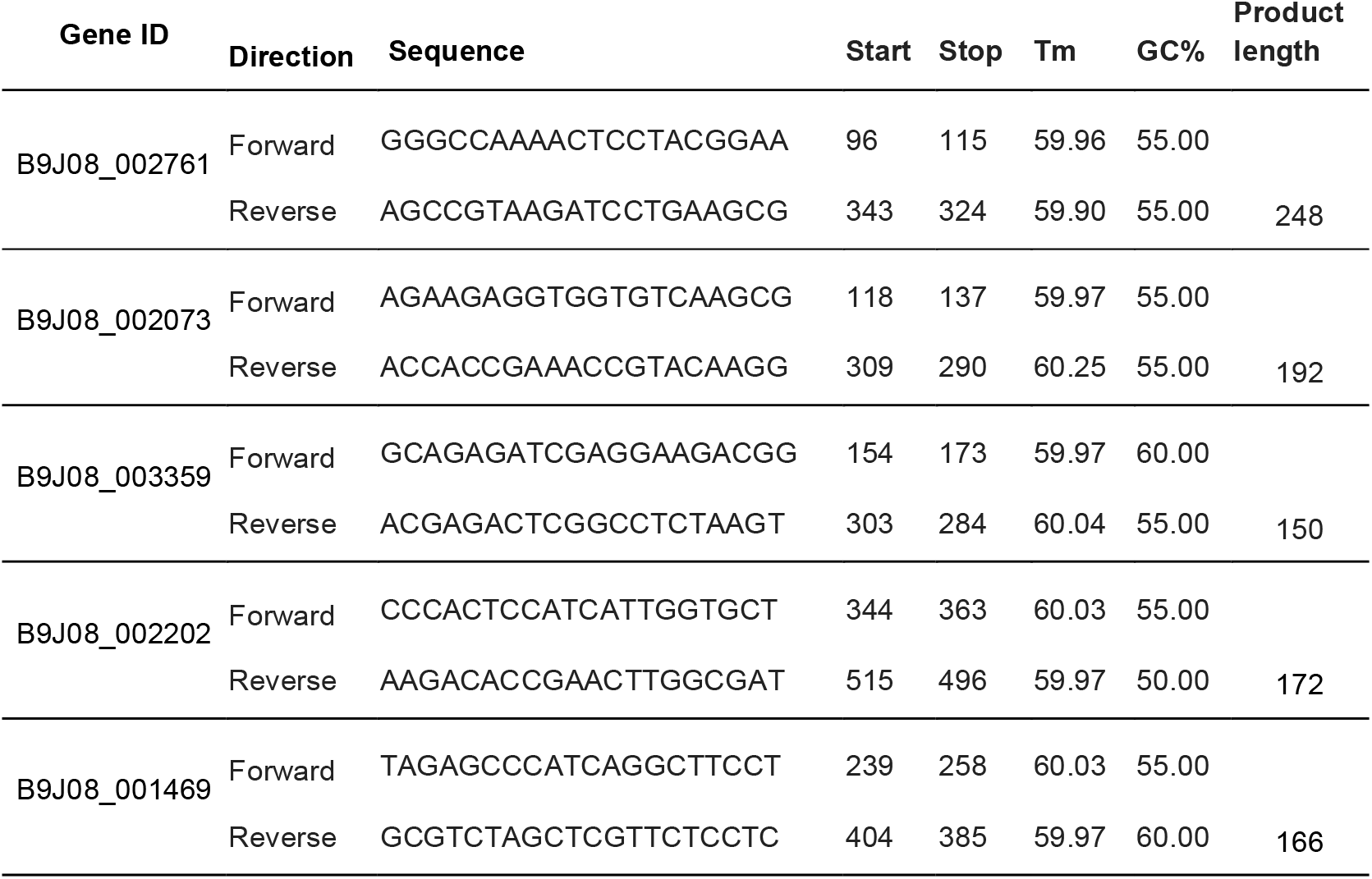
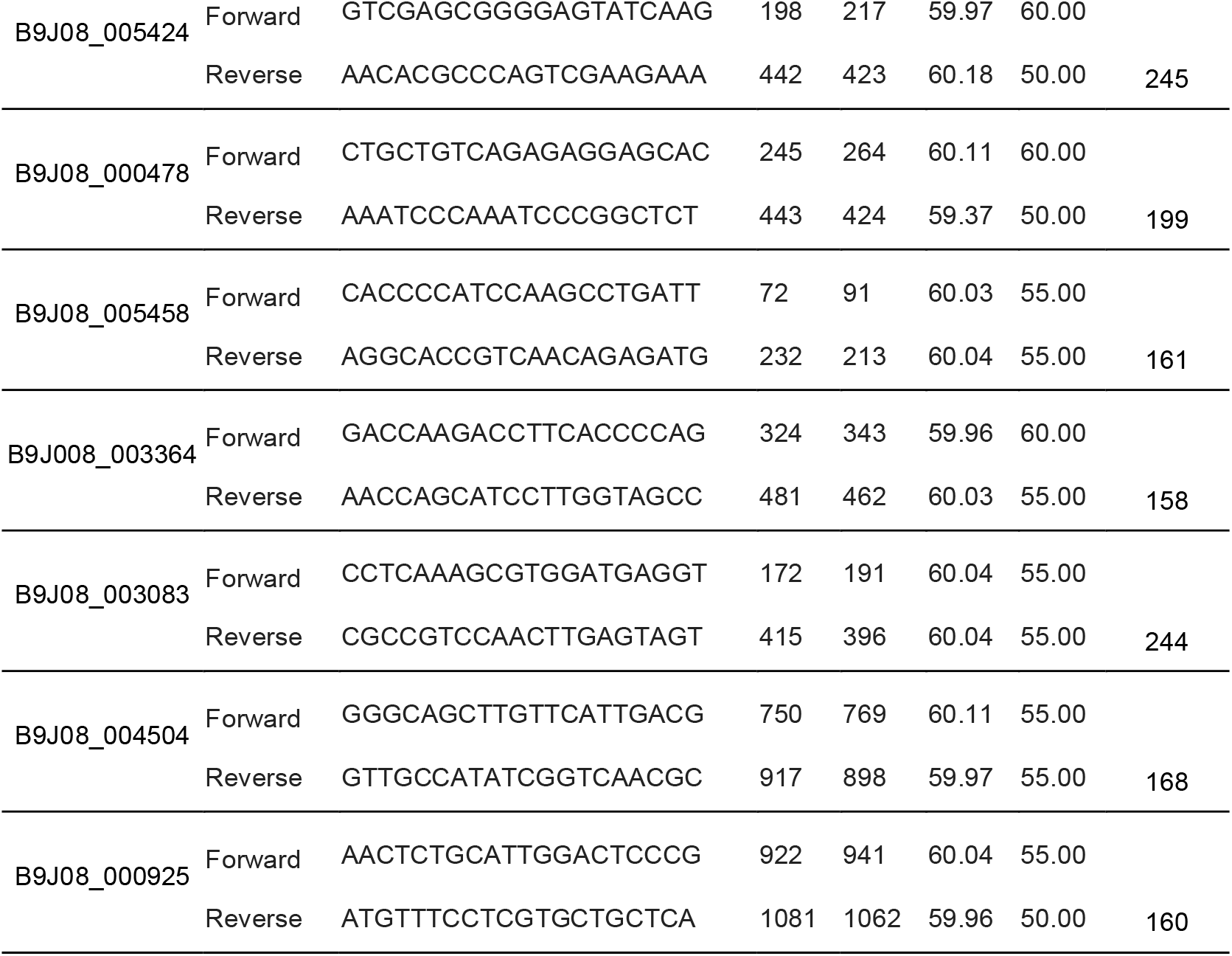
Primers used in this study

### Data analysis

The sequences in fastq format were analyzed by CLC Genomics Workbench© v 20.0 (Qiagen), using the corresponding *C. auris* genome for strain B8441 (GCA_002759435.2 V2). The parameters used for the alignments were: mismatch cost (2), insertion cost (3), deletion cost (3), length fraction (0.8), and similarity fraction (0.8). Only uniquely mapped reads were considered in the analysis. The statistical test applied was the DGE (Differential Gene Expression) using the RNA-seq package with CLC Genomics Workbench© v 20.0 (Qiagen). The expression values for the transcripts were registered in TPM (Transcripts per Million) and TMM (trimmed mean of M values) was used as a normalization method (34). The parameters to select the differentially expressed transcripts was 3-fold change (>3 FC) and False discovery rate (FDR) below or equal to 0.05.

The sequences generated from the small RNA-seq cell and EV were trimmed to remove the internal adapter that could be present in the shorter sequences. TruSeq small RNA adapter sequence: TGGAATTCTCGGGTGCCAAGG, and when the adapter was identified in the sequence, it was removed along with the following sequence (3’ trim).

For the ncRNA, the database used was the ncRNA from *Candida* genome databases and the reference available at the RNA central database (35): C_a uris_ B8441 _vers io n_sXX-mYY-rZZ_other_features_no_introns.fasta.gz and candida_auris_gca_001189475.ASM118947v1.ncrna. 1.

For the mRNA identification in the EVs, we combined the differential expression with reads coverage, so we performed the map reads to reference (C_auris_B8441_version_s01-m01-r10_genomic and C_auris_B8441_version_s01-m01-r10_other_features_plus_intergenic) using the following parameters: No masking, match score (1), mismatch cost (2), linear insertion cost (3), deletion cost (3), length fraction (0.6), similarity fraction (0.8) and global alignment. To consider the full-length mRNAs we selected those with expression value (TPM) higher than 100 and also 5x transcript coverage.

The differentially expressed transcripts sequence from MMC1 and B8441 cells were compared with orthologous genes from C. albicans genome strain SC5314 (assembly ASM18296v3) by Reciprocal Best Hit (RBH). The genes that presented more than 30% of identity were considered for the analysis. The Gene Ontology terms were generated using DAVID 6.8 software (36) and the String for protein interaction network and enrichment analysis (37). We applied the Fisher’s exact test and considered only the terms with p ≤ 0.05 and, next, we compared terms for the up- and down-regulated genes to a background of all terms to obtain an overall insight into the effect of caspofungin on C. auris compared to the absence of the antifungal.

## RESULTS

### Effects on C. auris induced by caspofungin

We studied two clinical isolates of *C. auris* with distinct caspofungin susceptibility profiles. *C. auris* strains B8441 and MMC1 were cultivated for 24 h in the absence or presence of caspofungin. The caspofungin concentration used for each strain was set to be strong enough to induce a stress response, but also low enough to allow fungal growth and EV production (Figure 1A, B). We examined if different caspofungin concentrations would elicit distinct morphological alterations in each *C. auris* strain. In untreated cells, well-defined, elliptical-shape yeast morphology as well as budding cells with typical bud scars were observed (Figure 1C, E). In contrast, cells treated with caspofungin exhibited a severely distorted yeast cell topography, with cells fused together and enlarged yeasts forming clumps mixed with cells with normal morphology (Figure 1D, F).

**Figure 1.**
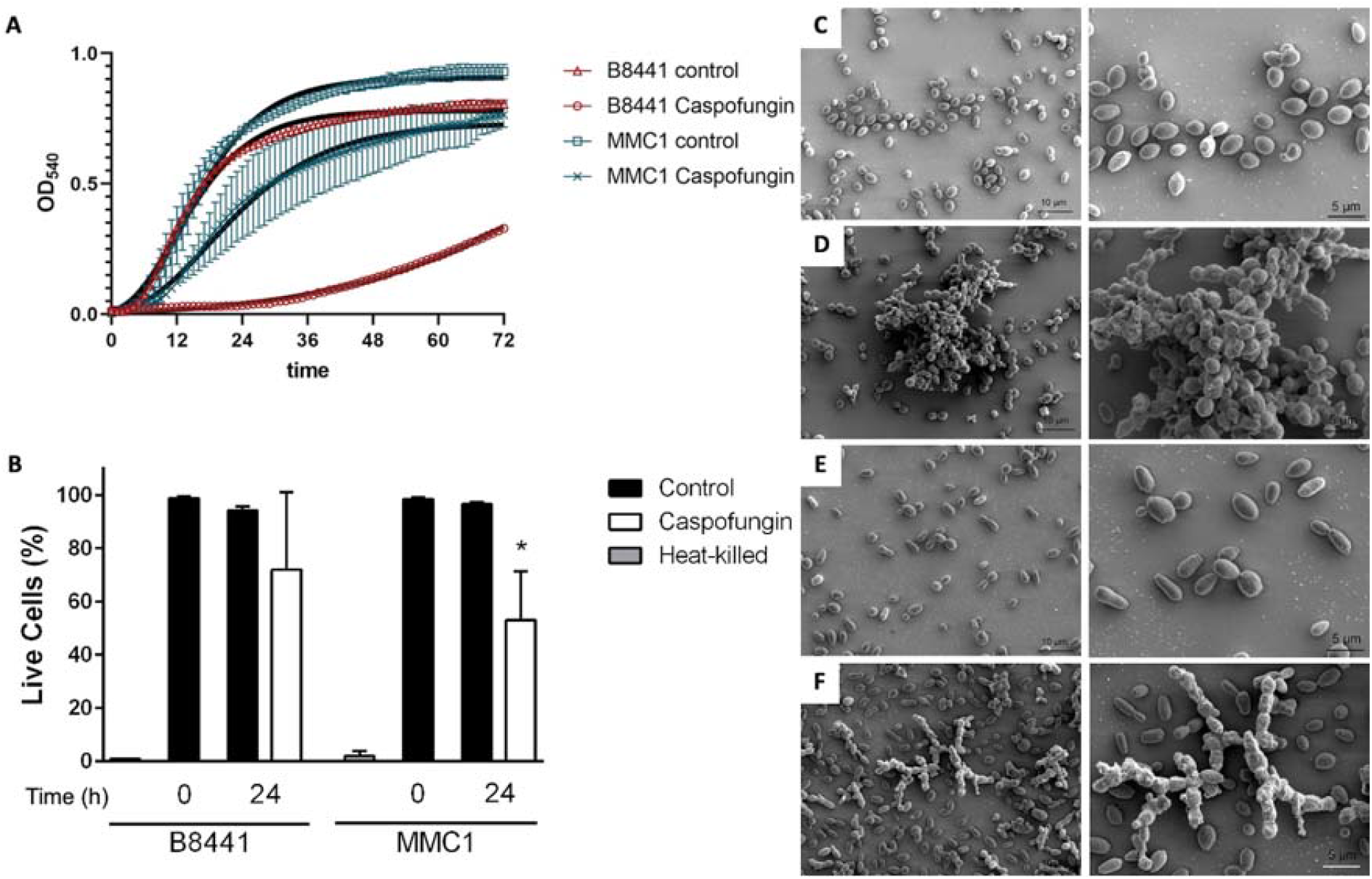
Caspofungin treatment led to morphological alterations in *C. auris*. Growth curve of *C. auris* in the presence of caspofungin (A). The growth was analyzed for MMC1 and B8441 strains and the assay was performed in a technical and biological triplicate. The significance was calculated by ANOVA. Effect of sub-optimal concentration of caspofungin on viability of *C. auris* (B). Cultures of *C. auris* were cultivated with or without caspofungin for 24 hours and cell viability was addressed by propidium iodide staining and analysis by flow cytometry. Graphs represent average and standard deviation for 4 independent experiments. * indicates p < 0.05 by paired t-test. Scanning electron microscopy images of *C. auris* at 1400x and 3000x magnification. Strain MMC1, control (C). MMC1 in Sabouraud with caspofungin (D) Strain B8441, control (E). Strain B8441 in Sabouraud with caspofungin (F).

As caspofungin is a drug that interferes with cell wall dynamics, we addressed important constituents of the cell wall after the treatment with caspofungin. Indeed, treatment with caspofungin induced modifications on the yeast cell wall of both strains of *C. auris*, as shown in Figure 1. Yeast cells treated with caspofungin had a thickened mannoprotein layer, as observed by microscopy (Figure 2A) and quantified by flow cytometry (Figure 2B). Other cell wall components such as chitin and chitin oligomers seemed increased by visual assessment; however, microscopy-based quantification is not accurate or sufficient to specifically confirm this finding. The stress response induced by caspofungin upon *C. auris* included an oxidative burst in both strains (Figure 2C). Caspofungin treatment interferes with the cell wall dynamics in *C. auris* as part of its mechanism of action and impacts the mannoprotein layer and the redox state of the treated yeast cells.

**Figure 2.**
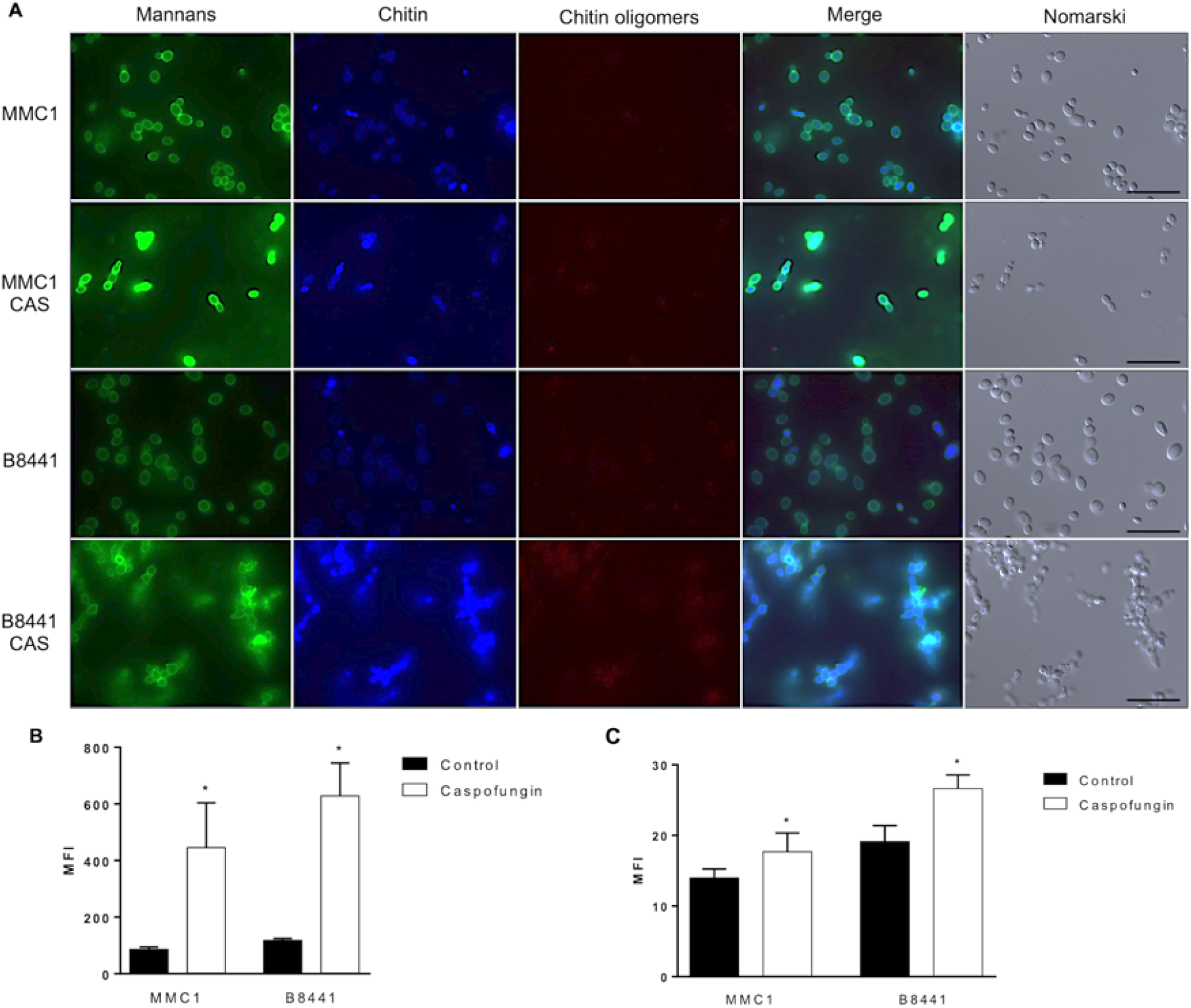
Effect of caspofungin on *C. auris* yeast cells. Yeast cells were treated with sub-lethal concentrations of caspofungin for 24 hours until the evaluation of cell wall components by microscopy (A) and mannoproteins by flow cytometry (B) or the production of reactive oxygen species by flow cytometry (C). * indicates *p* < 0.05 by one-way Anova followed by Bonferroni’s multicomparisons test for four independent experiments. Bars = 20 μm. CAS – caspofungin.

### Caspofungin treatment induces dynamic changes of gene expression in specific cellular processes

We investigated the *C. auris* gene expression changes in response to caspofungin treatment using RNA seq. *C. auris* strains B8441 and MMC1 were cultivated in three independent experiments for 24 h in the absence or presence of sub-inhibitory concentrations of caspofungin, and analyzed by RNA seq. We obtained on average 7.3 million reads per sample that mapped more than 95% of the reference genome (GCA_002759435.2), with a 30x coverage (Supplementary Table S1). We used Principal-component analysis (PCA) and hierarchical clustering to analyze the similarities between the triplicates and the differences among the treated and untreated samples. The triplicates clustered together, indicating a high level of correlation, whereas the treated and untreated samples clustered separately, consistent with a specific and global transcriptome response (Supplementary Figure S1).

To evaluate whether the transcripts were differentially expressed, we set the statistical significance of false-discovery rate (FDR) smaller than 5% and a fold change of 2 were considered a transcript differentially expressed. We found that 2085 transcripts were differentially expressed in the B8441 strain and 1588 in the MMC1 strain (Supplementary Table S2). When we compared the transcripts common to both strains, we observed 997 mRNAs, which corresponds to 47% and 64% of the differentially expressed transcripts in B8441 and MMC1 strains, respectively (Supplementary Table S3).

To evaluate the impact of caspofungin treatment in the gene expression of *C. auris*, we performed a functional enrichment analysis of the differentially expressed transcripts regulated in the presence of caspofungin in both MMC1 and B8441 strains. Firstly, we explored the 495 transcripts upregulated in the presence of the antifungal. Caspofungin induced changes in many important pathways such as cell cycle, nucleosome and various types of N-glycan biosynthesis (Figure 3). The most expressed transcripts in response to the antifungal were histones and GPI-anchored proteins (Table 2). Other significantly upregulated mRNAs in the biological process category were associated with cell wall biogenesis and protein acetylation, whereas genes in the molecular function category were related to nucleotide/GTP-binding and for the cellular component category there were upregulated genes involved with nucleosome and chromosome structures (Table 2 and Figure 3). Among the downregulated transcripts, we identified fewer pathways or processes enriched. In fact, regulation of transcription was the enriched term for biological processes and nucleus was the enriched term for cellular component (Table 3).

**Figure 3.**
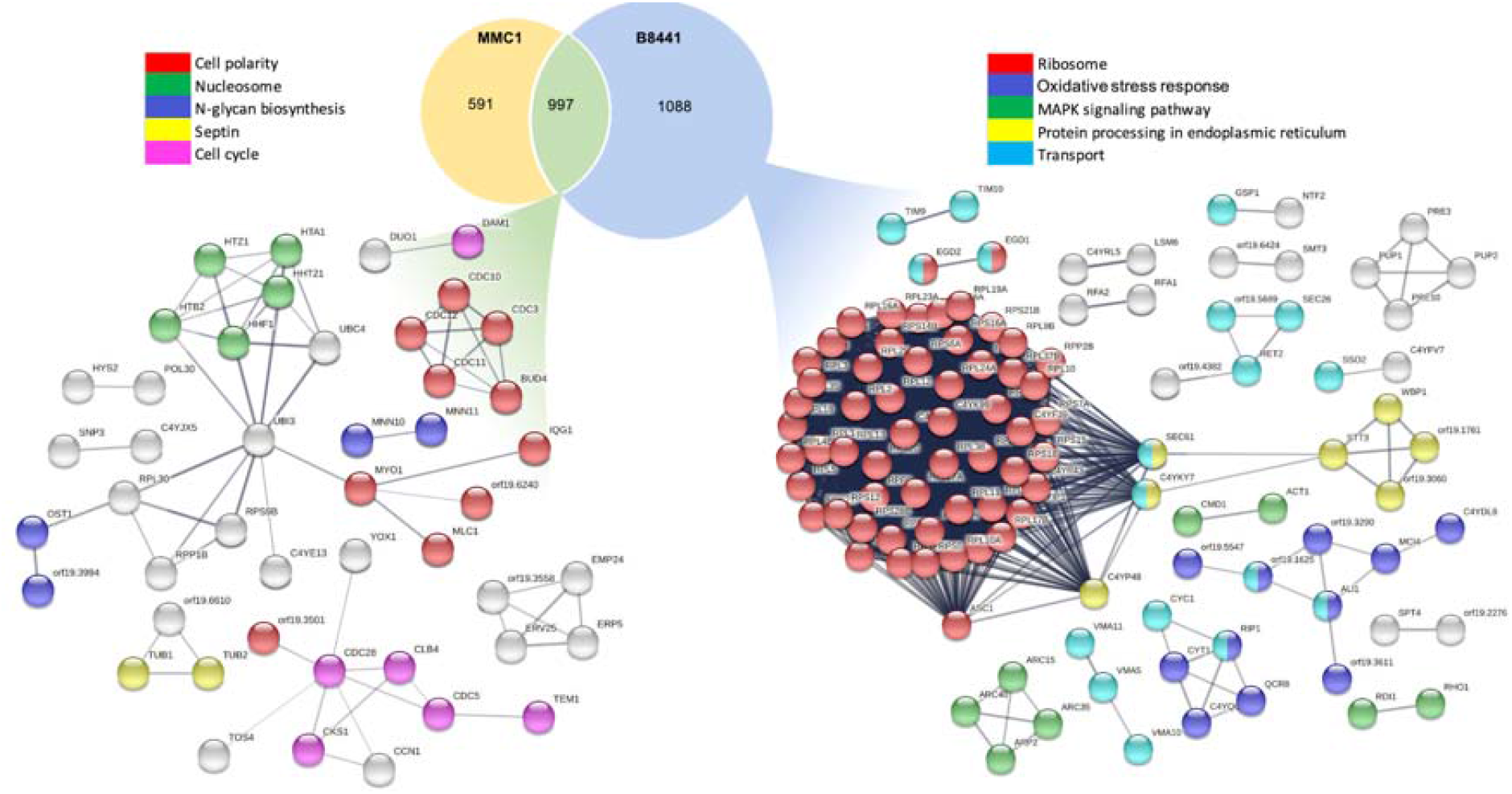
Enriched functional categorization of transcripts from *C. auris* strains MMC1 and B8441 yeast cells based on gene ontology (GO) annotations upon caspofungin treatment. B8441 n= 2085, MMC1 n= 1588, mRNAs identified in both strains n= 997. FDR ≤ 5%. For the analysis, only the transcripts with similarity with *C. albicans* greater than 40% were considered for the STRING analysis. On the left are the clusters and pathways exclusive to B8441 strain. On the right the clusters and pathways shared by both strains. At the top are the gene ontology terms.

**Table 2.** Most expressed upregulated and downregulated transcripts during caspofungin treatment.

|  | Name | Description | B8441 -<br>TPM<br>mean | B8441<br>- Log<sub>2</sub><br>fold<br>change | B8441 -<br>FDR p-value | MMC1 -<br>TPM<br>mean | MMC1 -<br>Log<sub>2</sub><br>fold<br>change | MMC1<br>- FDR<br>p-value | C. albicans<br>Uniprot ID |
| --- | --- | --- | --- | --- | --- | --- | --- | --- | --- |
| Upregulated | B9J08_001366 | Predicted GPI-anchored protein 6 | 4813.77 | 6.37 | 0.00% | 3943.45 | 6.57 | 0.00% | Q5APJ9 |
|  | B9J08_002073 | Histone H4 | 3939.53 | 6.19 | 0.00% | 5013.68 | 6.74 | 0.00% | Q59VN4 |
|  | B9J08_004382 | Cell surface GPI-anchored protein ECM33 | 858.66 | 5.94 | 0.00% | 1160.91 | 4.68 | 0.00% | Q5AGC4 |
|  | B9J08_002072 | Histone H3.1/H3.2 | 3913.19 | 5.92 | 0.00% | 4880.68 | 6.42 | 0.00% | Q59VN2 |
|  | B9J08_004381 | UDP-N-acetylglucamine diphosphorylase | 731.62 | 4.26 | 0.00% | 417.85 | 2.09 | 0.00% | A0A1D8PNG6 |
|  | B9J08_000438 | GPI-anchored protein 13 | 957.97 | 4.23 | 0.00% | 2422.00 | 2.29 | 0.00% | Q5A343 |
|  | B9J08_002600 | Phosphomannomutase | 780.86 | 4.13 | 0.00% | 906.33 | 2.74 | 0.00% | P31353 |
|  | B9J08_001462 | G1/S-specific cyclin CCN1 | 118.90 | 4.11 | 0.00% | 206.28 | 3.77 | 0.00% | Q59YH3 |
|  | B9J08_000354 | Mlc1p | 713.68 | 3.99 | 0.00% | 544.78 | 3.24 | 0.00% | A0A1D8PSE1 |
|  | B9J08_005437 | Probable pathogenesis-related protein | 1263.61 | 3.99 | 0.00% | 1364.49 | 4.62 | 0.00% | Q59WG5 |
|  | B9J08_005359 | Secreted beta-glucosidase SUN41 | 1435.47 | 3.94 | 0.00% | 2138.49 | 4.04 | 0.00% | Q59NP5 |
| Downregulated | B9J08_002202 | Phosphate transporter - PHO89 | 799.16 | -8.24 | 0.00% | 151.49 | -6.19 | 0.00% | Q5AMP8 |
|  | B9J08_001464 | Phosphate transporter - PHO84 | 1320.17 | -7.03 | 0.00% | 2046.23 | -3.49 | 0.00% | A0A1D8PF54 |
|  | B9J08_002660 | Hgt17p | 141.16 | -6.91 | 0.00% | 607.27 | -4.20 | 0.00% | A0A1D8PL74 |
|  | B9J08_004204 | Jen2p | 147.95 | -4.94 | 0.00% | 110.90 | -2.37 | 0.00% | A0A1D8PLY7 |
|  | B9J08_003046 | Cha1p | 118.12 | -4.38 | 0.00% | 223.30 | -1.82 | 0.00% | A0A1D8PGA8 |
|  | B9J08_001613 | Pho100p | 316.88 | -4.07 | 0.00% | 864.21 | -3.11 | 0.00% | Q59WE5 |
|  | B9J08_003409 | Transcriptional regulator of yeast form adherence 6 | 156.22 | -3.85 | 0.00% | 230.90 | -3.75 | 0.00% | Q5ADL8 |
|  | B9J08_003627 | Small heat shock protein 21 | 10549.82 | -3.67 | 0.00% | 8478.55 | -2.22 | 0.00% | Q5AHH4 |
|  | B9J08_001012 | SWIRM domain-containing protein | 978.15 | -3.62 | 0.00% | 276.07 | -1.86 | 0.00% | A0A1D8PII2 |
|  | B9J08_003998 | Acetyl-coenzyme A synthetase | 779.18 | -3.58 | 0.00% | 2764.50 | -3.46 | 0.00% | Q59XW4 |
|  | B9J08_002840 | Stress-responsive transcriptional activator | 444.33 | -3.53 | 0.00% | 847.06 | -2.75 | 0.00% | A0A1D8PEH3 |
|  | B9J08_001679 | Transcription factor CPH2 | 1046.33 | -2.23 | 0.00% | 1035.95 | -1.74 | 0.00% | Q59RL7 |

**Table 3.** Gene ontology terms for the downregulated transcripts in the presence of caspofungin.

**Gene ontology - mRNAs downregulated B8441 and MMC1**
| Term | Count | % | P-Value | Fold Enrichment |
| --- | --- | --- | --- | --- |
| nucleus | 13 | 13.7 | 1.6E-2 | 2.1 |
| amino acid transporter | 3 | 3.2 | 2.8E-2 | 9.3 |
| Transcription regulation | 7 | 7.4 | 1.4E-2 | 3.5 |

**Gene ontology - mRNAs downregulated exclusive B8441**
| Term | Count | % | P-Value | Fold Enrichment |
| --- | --- | --- | --- | --- |
| DNA-binding | 13 | 6.8 | 4.9E-5 | 4.3 |
| Tyrosine metabolism | 5 | 2.6 | 6.5E-4 | 11.8 |
| nucleus | 27 | 14.1 | 8.7E-4 | 1.9 |
| Transcription | 13 | 6.8 | 1.8E-3 | 2.9 |

We then analyzed the transcripts exclusively expressed by B8441 where 528 transcripts were upregulated in the presence of caspofungin. The enriched pathways associated with these mRNAs were ribosome, oxidative stress response, phagosome and protein processing in endoplasmic reticulum (Figure 3). The most differentially expressed mRNAs were associated with ribosomal proteins, oxidative response to stress and protein transport (Supplementary Table S3). The 560 downregulated transcripts upon caspofungin treatment in the B8441 strain included numerous mRNAs coding for transcription factors, nucleus and DNA binding (Table 2). Notably, the increased transcript levels correlated with protein abundance. For example, histone 4 and ribosomal protein S6 were more expressed in the cells treated with caspofungin, reinforcing our observation at the transcript level (Supplementary Figure S2).

### Characterization of EVs produced by C. auris treated with caspofungin

Next, we investigated whether caspofungin could affect the extracellular vesicles (EVs) RNA cargo in *C. auris*. EVs were isolated from the supernatant of *C. auris* cultures treated or not with caspofungin. The EVs displayed the standard morphology of “cup-shaped” lipid bilayered vesicles of typical sizes (Figure 4A-D) as previously described (38). There were no differences in the morphology in EVs from cultures treated or not with caspofungin, indicating that the detection of EV-like particles in the drug-treated systems was not a consequence of the leakage of intracellular organelles. The amount of EVs isolated doubled from cultures subjected to caspofungin treatment (Figure 4E). Nano tracking particle analysis revealed that *C. auris* EVs isolated from MMC1 and B8441 strains presented a slight difference in size, as the EVs isolated in the presence of the antifungal tended to be bigger compared to the control (Figure 4F,G). Caspofungin-treated *C. auris* produced 2 times more EVs than non-treated *C. auris*. EV morphology, however, was not substantially changed because of caspofungin.

**Figure 4.**
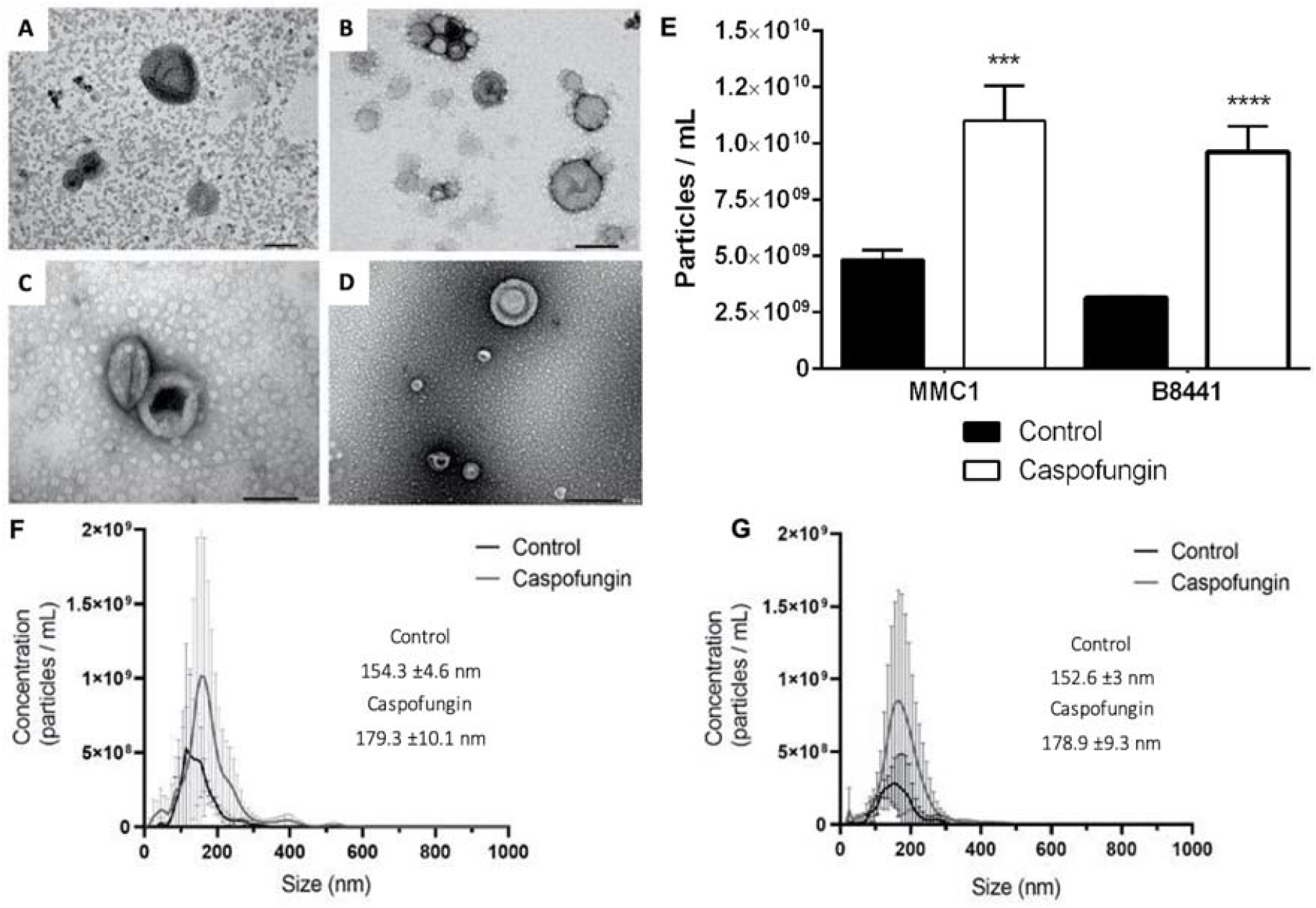
Characterization of *C. auris* extracellular vesicles (EVs). Transmission electron microscopy images of MMC1 strain EVs isolated from control conditions (A) caspofungin treatment (B). Transmission electron microscopy of B8441 strain EVs derived from control conditions (C) or caspofungin treatment (D), bar = 100 nm. Bar chart representing EVs concentration calculation in both strains from control and caspofungin treatment (E), the X-axis indicate the particles concentration, *** p < 0.001 and **** p < 0.0001, by one-way Anova followed by Bonferroni’s multicomparisons test for three independent experiments. Representative comparison size distribution graphs for MMC1 (F) EVs and B8441 (G) from caspofungin (grey line) and control (black line), respectively. The average EV size is indicated in nm.

### Caspofungin led to alteration in the composition of RNA in EVs

In addition to exploring the alterations caused by caspofungin at the cellular level, we hypothesized that the RNAs directed to the EVs would also be affected. Indeed, for EVs-RNA, there was a difference in composition when the cells were treated with caspofungin. We identified distinct RNA molecules as full-length mRNAs; however, the bulk was composed by molecules smaller than 200 nucleotides that included fragments of mRNAs, tRNAs, other ncRNAs and rRNA, as previously described in other fungi species (39). It is noteworthy that when we extracted RNA from B8441 EVs isolated from cells cultured in the presence of caspofungin, the yield of vesicular RNA was 10 x higher than MMC1 EVs in the same condition, and this is reflected on the analysis, where we obtained a distinct profile of RNA molecules comparing B8441 and MMC1 (Supplementary Table S4).

The library construction for small RNA is based on size selection of the fragments. Therefore, for EV library preparation, we selected not only the small fraction of RNAs but also the longer molecules, that correspond to the mRNAs present in the vesicles. After sequencing, to discriminate full-length mRNAs, we selected transcripts with a 3x of minimum reads coverage and expression value (expressed in transcripts per million) greater than 100. Applying these filters, 166 transcripts abundant in the B8441 strain and 31 in the MMC1 strain were identified (Supplementary Table S4).

For the B8441 strain, we observed 20 full length transcripts contained in the EVs isolated from yeast cultured in standard media, and the most abundant were omega-class glutathione transferase, phosphatidylinositol-3-phosphate-binding, ubiquitin-protein ligase, aminodeoxychorismate lyase, E1 ubiquitin-activating protein among others (Table 3). However, caspofungin induced alterations in 146 of the identified transcripts. There was an enrichment for the cell wall, nucleosome core and ribosome, and the latter is similar to what was observed for the transcriptome when cells were treated with caspofungin (Figure 5A). Among the transcripts present in EVs from B8441 and MMC1, 13 were common, reinforcing the result observed for the transcriptome indicating that *C. auris* responded to the antifungal in a common way. We also observed fragments of mRNAs in the EVs, and most of them were related to mitochondrial metabolism in the presence of caspofungin (Table 4).

**Figure 5.**
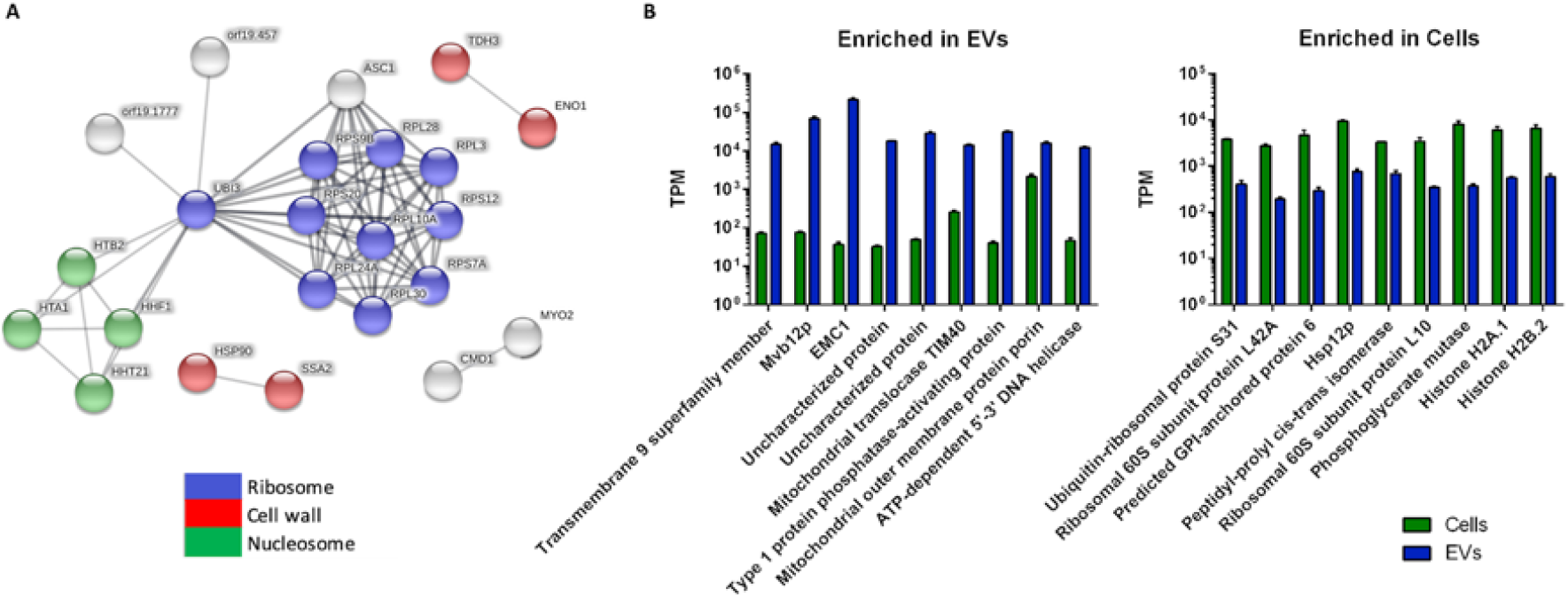
Gene ontology analysis of full-length mRNA identified in *C. auris* EV isolated from B8441 strain yeast cells treated with caspofungin (A). * FDR ≤ 5%. Comparison between mRNAs enriched in the EVs compared to the cell (B).

**Table 4.** Full-length mRNAs present in the EVs.

|  | Reference ID | COVERAGE<br>Control | COVERAGE<br>Caspo | Uniprot C.<br>albicans | Description | TPM<br>mean | Log<sub>2</sub><br>fold<br>change | FDR |
| --- | --- | --- | --- | --- | --- | --- | --- | --- |
| mRNAs enriched in EVs - Caspofungin | **B9J08_002056** | **187.16** | **5298.89** | **A0A1D8PN56** | **EMC1** | **91990.36** | **4.12** | **0.0%** |
|  | **B9J08_003626** | **83.93** | **605.97** | **Q5AHH6** | **Type 1 protein phosphatase-activating protein** | **21139.63** | **2.38** | **0.2%** |
|  | **B9J08_001980** | **23.12** | **522.25** | **A0A1D8PSU3** | **Mvb12p** | **31832.19** | **4.17** | **0.0%** |
|  | B9J08_001788 | 12.26 | 291.84 | Q5AF62 | Transmembrane 9 superfamily member | 7489.03 | 3.26 | 0.0% |
|  | **B9J08_005458** | **19.41** | **249.89** | **P83774** | **Guanine nucleotide-binding protein subunit beta-like protein** | **2678.05** | **3.45** | **0.0%** |
|  | B9J08_000654 | 25.08 | 150.36 | Q5ABA2 | Survival factor 1 | 3131.41 | 2.33 | 0.5% |
|  | B9J08_004879 | 13.14 | 122.98 | A0A1D8PL12 | MHD domain-containing protein | 4752.14 | 2.93 | 0.0% |
|  | B9J08_002476 | 19.78 | 111.22 | Q5A201 | Transcriptional regulator GZF3 | 4862.05 | 1.94 | 1.7% |
|  | **B9J08_005403** | **13.69** | **107.27** | **Q59XM0** | **MFS antiporter QDR3** | **3283.26** | **2.65** | **0.1%** |
|  | B9J08_003261 | 9.75 | 79.94 | O94030 | Mitochondrial intermembrane space import and assembly protein 40 | 12954.17 | 3.35 | 0.0% |
| mRNAs enriched in EVs - Control | B9J08_001103 | 42.96 | 0.55 | A0A1D8PS56 | Omega-class glutathione transferase | 2140.41 | -2.52 | 1.7% |
|  | B9J08_003689 | 41.58 | 0.75 | A0A1D8PU28 | Phosphatidylinositol-3-phosphate-binding ubiquitin-protein ligase | 4299.11 | -2.95 | 0.6% |
|  | **B9J08_003083** | **28.54** | **0.83** | **Q5A0H4** | **Aminodeoxychorismate lyase** | **31254.92** | **-4.15** | **0.0%** |
|  | B9J08_001654 | 24.75 | 0.98 | A0A1D8PSA4 | E1 ubiquitin-activating protein | 4594.68 | -1.92 | 5.4% |
|  | B9J08_000375 | 11.73 | 1.25 | Q59KQ3 | Uncharacterized protein | 1745.48 | -1.98 | 1.1% |
|  | B9J08_000925 | 10.75 | 0.28 | Q5A1Z5 | Autophagy-related protein 13 | 1073.19 | -3.14 | 0.4% |
|  | B9J08_004718 | 10.24 | 1.21 | A0A1D8PE79 | Oxysterol-binding protein related protein | 801.93 | -2.39 | 0.2% |
|  | B9J08_005450 | 9.63 | 0.77 | Q5AKU6 | Oxidative stress response two-component system protein SSK1 | 931.46 | -2.25 | 0.8% |
|  | **B9J08_002986** | **6.27** | **0.26** | **A0A1D8PQ59** | **1,4-alpha-glucan branching enzyme** | **950.02** | **-2.63** | **0.5%** |
|  | B9J08_004711 | 5.48 | 1.29 | Q5A2I3 | RING-type domain-containing protein | 667.83 | -1.86 | 0.2% |

In bold are the mRNAs that were also present in the MMC1 strain.

We next checked whether the most expressed mRNAs in the cell were also present in the EVs of the B8441 strain (Figure 5B). The high level of expression of a determined transcript did not reflect the presence of the RNA in the EVs. In fact, the full-length mRNAs in the EVs presented average levels of expression when the whole transcriptome was taken into consideration (Figure 5B).

To validate the data from our RNA-Seq analysis from both cell and EVs, we performed real-time quantitative PCR (qPCR) on MMC1 and B8441 strains under control condition and with caspofungin treatment. We selected 3 genes with highly abundant transcripts in EVs as well as the 3 of the most expressed genes in the cell. For all selected targets, qPCR results were in accordance with the results from the vesicular and cellular RNA-seq analysis (Supplementary Figure S3).

Next, we wanted to investigate if the differences in EV RNA composition compared to the cell could be due to the differences in the library construction rather than in biological processes, which could lead to a biased interpretation of the results. To test this hypothesis, we constructed a library with the small fraction of the cellular RNA (sRNA) and also performed the size selection within the same range as for the EVs. In fact, we did not observe an enrichment of specific full-length mRNAs differentially abundant in the sRNA cell content as we observed for the EV mRNAs (Supplementary Table S5).

Another class of RNA molecule represented in the EVs were the ncRNAs. For B8441 strain, 109 ncRNAs were identified with 33 being differentially abundant during caspofungin treatment, 45 in the control and 29 equally identified in both conditions (Supplementary Table S6). When comparing the two strains, 49 identified ncRNAs were common. From the identified ncRNAs, tRNAs were the most prevalent molecules in B8441 and MMC1. Most of the tRNAs identified were tRNA-derived fragments (tfRNAs).

The ncRNA content of the cell was also analyzed. For control conditions in both strains, we detected tRNAs and snoRNAs as the most prevalent molecules. Regarding tRNAs, we could observe complete tRNAs as well as tfRNAs. However, there was a shift when the cells were treated with caspofungin, and rRNA became the most identified ncRNA molecule in both strains (Figure 6A). Despite the greater number of rRNA identified in the cells treated with caspofungin, this did not reflect their identification in the EVs released in the presence of the antifungal (Figure 6B).

**Figure 6.**
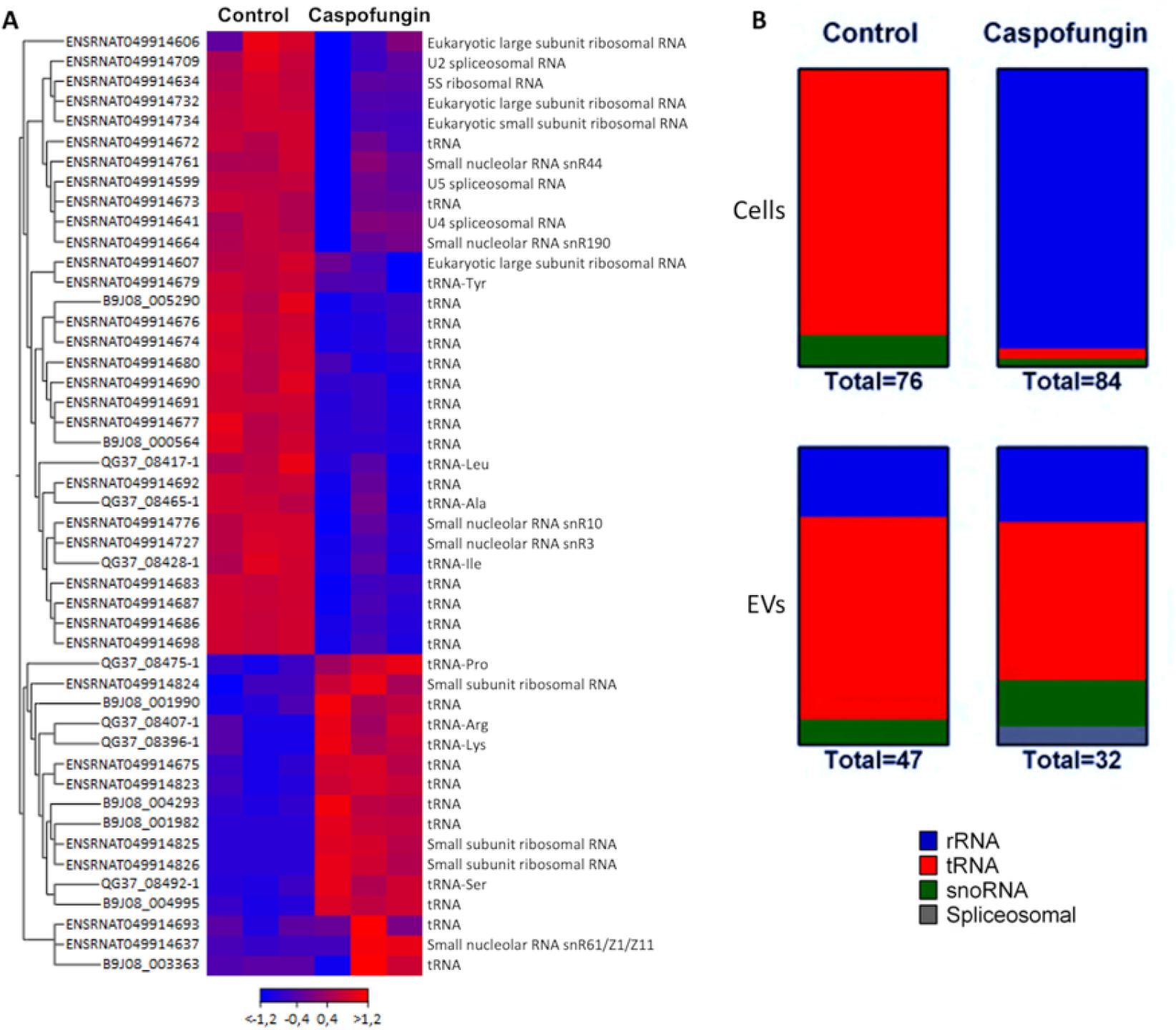
ncRNAs analysis. (A) Heat map representing expression levels of differentially abundant ncRNAs present in EVs for B8441 strain comparing control vs. caspofungin (FDR <5% and fold change up- or down-regulated >4-fold). The expression levels are visualized using a gradient color scheme, blue – high expression level, red – low expression level. (B) Distribution of ncRNAs in *C. auris* yeast cells and EVs under control conditions and caspofungin treatment. The values refer to relative expression of the distinct classes of ncRNAs.

## DISCUSSION

Caspofungin treatment led to alterations in *C. auris* cell morphology and increased EVs release. In the presence of this drug, aberrant morphologies have been observed in other studies of *Candida* species (40, 41), including *C. auris*, when the cells were treated with a novel 1,3-β-D-glucan synthesis inhibitor (42). In addition, caspofungin treatment increases EV production by *S. cerevisiae* (32).

To gain novel insights into *C. auris* gene expression in response to echinocandins, we performed a transcriptomic analysis of B8441 and MMC1 strains grown in the presence or absence of caspofungin. Concerning the transcripts with altered expression upon caspofungin treatment, we identified multiple pathways that were significantly modified. Molecules involved in N-glycan biosynthesis were enriched in both strains treated with caspofungin, which is consistent with cell wall rearrangement. In *C. neoformans*, N-linked glycosylation is an important modulator of host cell death, and therefore has a critical role in pathogenicity (43). Cryptococcal mutants harboring truncated N-glycans were not pathogenic in mice, despite being able to attach to lung epithelial cells and enter them through phagocytosis. Also, their capacity for cell wall remodeling was maintained, but they induced less cell death in macrophages, a mechanism used for pulmonary escape and dissemination *in vivo* (43).

Cell wall related transcripts, such as glycosylphosphatidylinositol (GPI)-anchored associated genes, were the most expressed genes in the presence of caspofungin for both strains. Interestingly, cell wall genes were previously shown to be upregulated during the initial hours of biofilm formation (19) (Supplementary Table S2). This high expression levels of cell wall related transcripts could be a compensatory mechanism to circumvent the absence of glucans with other components, such as chitin and mannans, which may reduce *C. auris* susceptibility to antifungal drugs. In fact, our results support such compensatory mechanism, evidenced by altered levels of transcripts, and mannans and chitin. This outcome has been described for several *Candida* species that rapidly responded to caspofungin by increasing the cell wall chitin content (44, 45). In addition, a similar result was observed in *A. fumigatus* where caspofungin treatment led to an increased chitin content, reduced susceptibility to the antifungal and altered cell morphology (46).

The mRNA coding GPI-anchored protein ECM33 was one of the most upregulated transcripts in both *C. auris* strains under caspofungin treatment. ECM33 is responsible for assembling the mannoprotein outer layer of the cell wall (47). Supporting our mRNA data, *C. auris* yeast cells from both strains presented a dramatic increase in their mannoprotein layer under caspofungin treatment.

Cell wall stressors like caspofungin induce an oxidative burst in fungi, including *C. albicans* (48, 49). Our data not only showed a caspofungin-induced upregulation of transcripts related to this pathway but also induced an increase in ROS production in both strains of *C. auris*. Similar observations were reported for *C. albicans*, of which caspofungin-induced cell wall stress increased ROS generation (50). Caspofungin also affects the oxidative stress response in *A. fumigatus*, of which many mRNAs related to this pathway are upregulated in the presence of the antifungal (51).

We also observed a higher expression of mRNAs coding for ribosomal proteins in both strains treated with caspofungin. These results are in agreement with the previous studies characterizing the *C. auris* transcriptome during amphotericin B and voriconazole treatment (52). Upon amphotericin B treatment, the upregulated genes were related to translation and ribosomal proteins (52). Compared to our data, we found 78 common transcripts also upregulated in the presence of caspofungin. From the shared mRNAs, most responded to amphotericin B treatment in which 46 were also identified in our data. We also found 30 transcripts upregulated during both caspofungin and voriconazole treatments. From these common transcripts, the majority of the mRNAs code for ribosomal proteins (Supplementary Table S2). In addition, during the biofilm formation in *C. auris*, the most consistent mRNAs upregulated during the process were those coding ribosomal proteins (19). In a study that combined proteomics and microarray analyses of *Aspergillus fumigatus* treated for 24 h with caspofungin, 81% of the overexpressed proteins and 86.4% of the mRNAs were ribosomal (53). This change was suggested to be associated with a ribosomal reshuffling response, which reflects a requirement for more protein synthesis to overcome the inhibition caused by the antifungal drug (54).

Our results demonstrated a high expression of genes related to nucleosome and DNA replication in caspofungin treated cells, as well as several mRNAs coding for histones. In *C. albicans*, the availability of specific chromatin modifiers can affect drug resistance (55). The transcription factor Cas5 has been implicated in stress responses, drug resistance and cell cycle regulation in *C. albicans* (56). In the presence of caspofungin, the stress caused by this antifungal led to distinct response, and the upregulated pathways were related to cell cycle, DNA replication and meiosis (56).

Next, we investigated how the response to caspofungin affected the RNA content of *C. auris* EVs. Our analysis demonstrated a low expression in cells of some highly enriched transcripts in EVs. This result indicates a regulated selection process for the molecules directed to the EVs, as previously described in mammalian and other fungal cells (57, 58). Both *C. auris* cells and their derived EVs had similar gene ontology terms but distinct enrichment of different terms. This allowed us to conclude that mRNAs are targeted to EVs in a specific mechanism, which is regulated by cell responses to a stimulus.

The most abundant transcript in the EVs after exposure to caspofungin was EMC1 (<u>e</u>ndoplasmic reticulum <u>m</u>embrane <u>c</u>omplex). EMC is a conserved protein complex of the ER that is necessary for homeostasis, and membrane protein biogenesis (59). EMC1 has been linked to antifungal resistance, as deletion of EMC1 led to an increased sensitivity to the sr7575 drug in *S. cerevisiae* (60). We hypothesized that this transcript could be transferred between cells upon caspofungin treatment, which is supported by our data showing that caspofungin-treated *C. auris* produces more EVs. Another example is the MVB12 (Multivesicular body sorting factor 12), an important component of ESCRT-I complex, required for cargo sorting into multivesicular bodies and EV biogenesis in other eukaryotes (61). Another mRNA enriched in EVs from caspofungin-treated cells is the type 1 protein phosphatase-activating protein (SDS22), which regulates the expression of pathogenic determinants in *C. albicans* (62). We speculate that the presence of this transcript inside of *C. auris* EVs is related to cell wall damage caused by caspofungin. The transcript coding for QDR3, a drug transport regulator, in EVs from caspofungin treated cells might suggest a communication in antifungal response. In *C. albicans*, QDR3 participates in the development of biofilm and virulence (63).

The transcript coding the MHD domain-containing protein was upregulated in EVs derived from the B8441 strain treated with caspofungin. This mRNA participates in septin cytoskeleton organization (64). In *C. albicans*, septin regulation is crucial in the early caspofungin response (65). Septins belong to the protein family of GTP-binding proteins that help define cell shape by contributing to cytokinesis, membrane remodeling, scaffolding, and chitin deposition in the cell wall (66). This result suggests that EV-mRNAs play a role in drug-induced stress signaling and cell-wall maintenance.

There is a paucity of studies discussing the role of ncRNAs in eukaryotic pathogens. In the parasites *Plasmodium falciparum* (67) and *Schistosoma* (68), virulence-related ncRNAs are expressed differentially according to the sexual stage of the organism, and this differential expression aids mechanisms such as the scape from host’s immune system and biotrophic survival during the infection process (67). In fungi, ncRNAs are also involved in the dimorphic aspect of cells, a process recognized as important in the context of infection and virulence (69). In our study, under normal growth conditions, the tRNA-halves were the prevalent ncRNA molecules identified in the EVs; however, for B8441 strain, treated with caspofungin, the ribosomal RNA was the prevalent ncRNA identified in the EVs. ncRNAs were enriched in the *C. auris* EVs, as similarly described for other eukaryotes. For example, *Paracoccidioides* had up to 71 different sequences of ncRNAs, with tRNA-fragments as the more prevalent molecules (70). This pattern was also observed in our previous studies of *C. albicans, C. neoformans* and *S. cerevisiae* (39). The biological significance of these findings is still unknown, but these results reinforce the idea that the fungal response to caspofungin affects several physiological processes.

In summary, our results show alterations in the distribution of RNA in EVs derived from yeast cells treated with an antifungal (Figure 7). The results showed that genes coding ribosomal proteins, cell wall, and cell cycle were significantly up-regulated in cells in the presence of caspofungin, whereas transcriptional regulation were down-regulated. Caspofungin also led to a transcriptional reduction of pathways related to oxidative stress. The upregulation of various types of N-glycan biosynthesis are related to cell wall biogenesis and the alterations in genes expression associated to wall synthesis could be a compensatory mechanism due to the blocking of β1,3-glucan synthesis by caspofungin. Altogether, these results from the transcriptome and EV RNAs provide further insights into the biological activity and mode of action for *C. auris* growth in the presence of caspofungin. This information expands our understanding of fungal drug resistance, a serious and emerging global threat.

**Figure 7.**
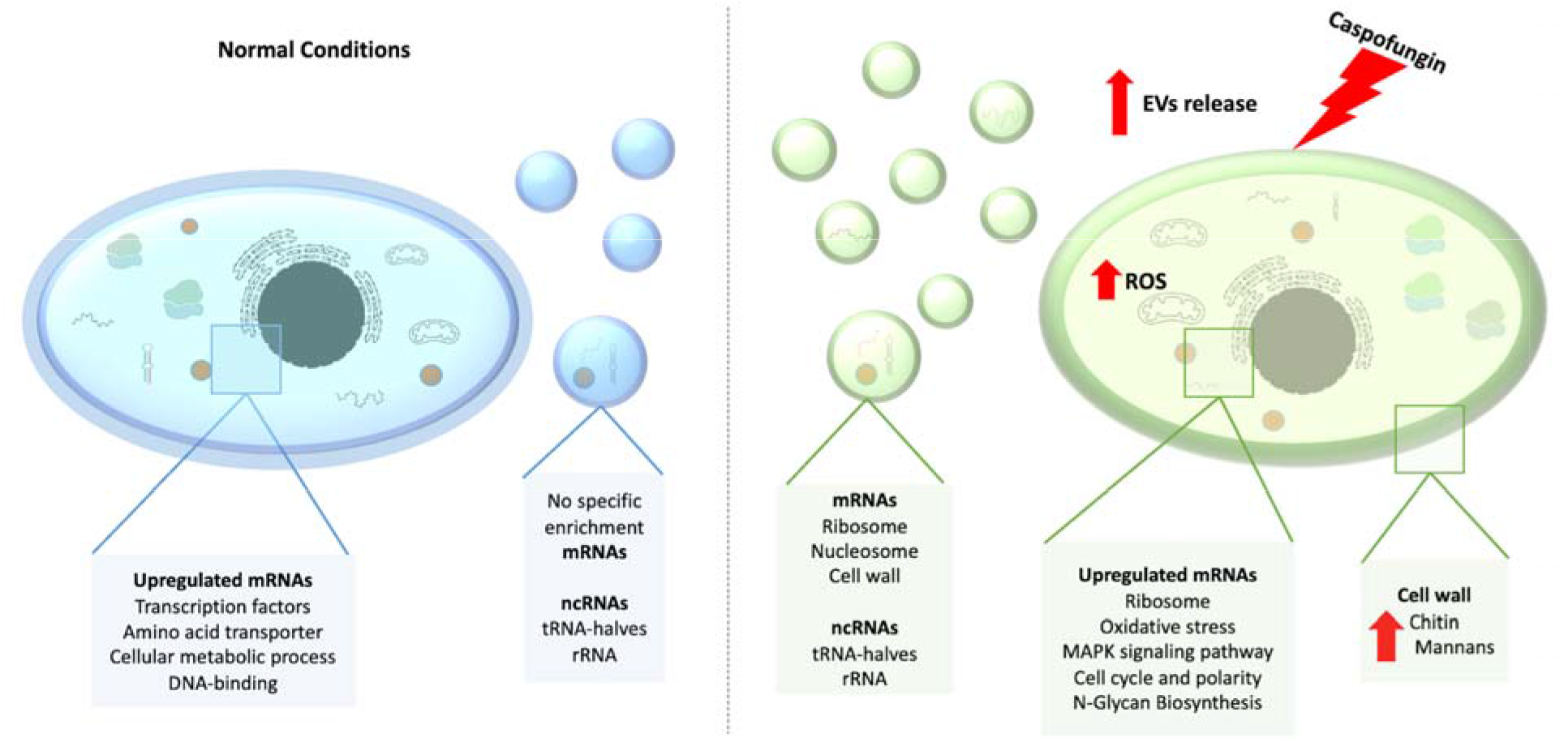
Schematic diagram of *C. auris* response upon caspofungin treatment.

## ACKNOWLEDGMENTS

We thank the staff of the Genomics section of the Life Sciences Core Facility (LaCTAD), part of the University of Campinas (UNICAMP), for their contributions to RNA-sequencing. J.D.N., D.Z-M. and E.S.N. were partially supported by NIH R21 AI124797. M.L.R. was supported by grants from the Brazilian Ministry of Health (grant number 440015/2018-9), Conselho Nacional de Desenvolvimento Científico e Tecnológico (CNPq, grants 405520/2018-2, and 301304/2017-3) and Fiocruz (grants VPPCB-007-FIO-18 and VPPIS-001-FIO18). The authors also acknowledge support from the Instituto Nacional de Ciência e Tecnologia de Inovação em Doenças de Populações Negligenciadas (INCT-IDPN). M.L.R. is currently on leave from the position of Associate Professor at the Microbiology Institute of the Federal University of Rio de Janeiro, Brazil. LRA received financial support from Inova Fiocruz/Fundação Oswaldo Cruz [Grant number VPPCB-07-FIO-18-2-52] and CNPq [Grant number 442317/2019-0]. L.R.A is a research fellow awardee from CNPq.

## DATA AVAILABILITY

The RNA-seq data have been deposited at the Sequence Read Archive (SRA) database under the accession number (SRA: SRP295539 BioProject: PRJNA682185).

## SUPPLEMENTARY DATA

Supplementary Data are available at NAR Online

