## Supplementary figures S1, S2 and S3 for "Integrated transcriptional analysis of the cellular and extracellular vesicle RNA content of Candida auris in response to caspofungin"

**SUPPLEMENTAL FIGURES**

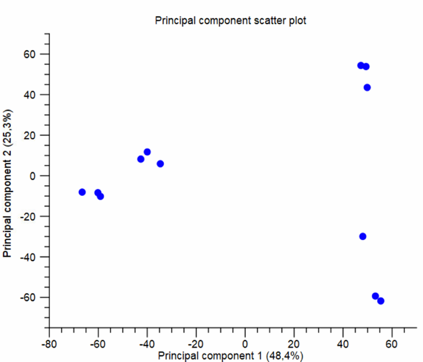

B8441

Caspofungin

B8441

MMC1

MMC1

Caspofungin

S

**Supplemental Figure 1**. RNA-seq experiment quality control. Principal component analysis evidencing the variation between samples and replicates.

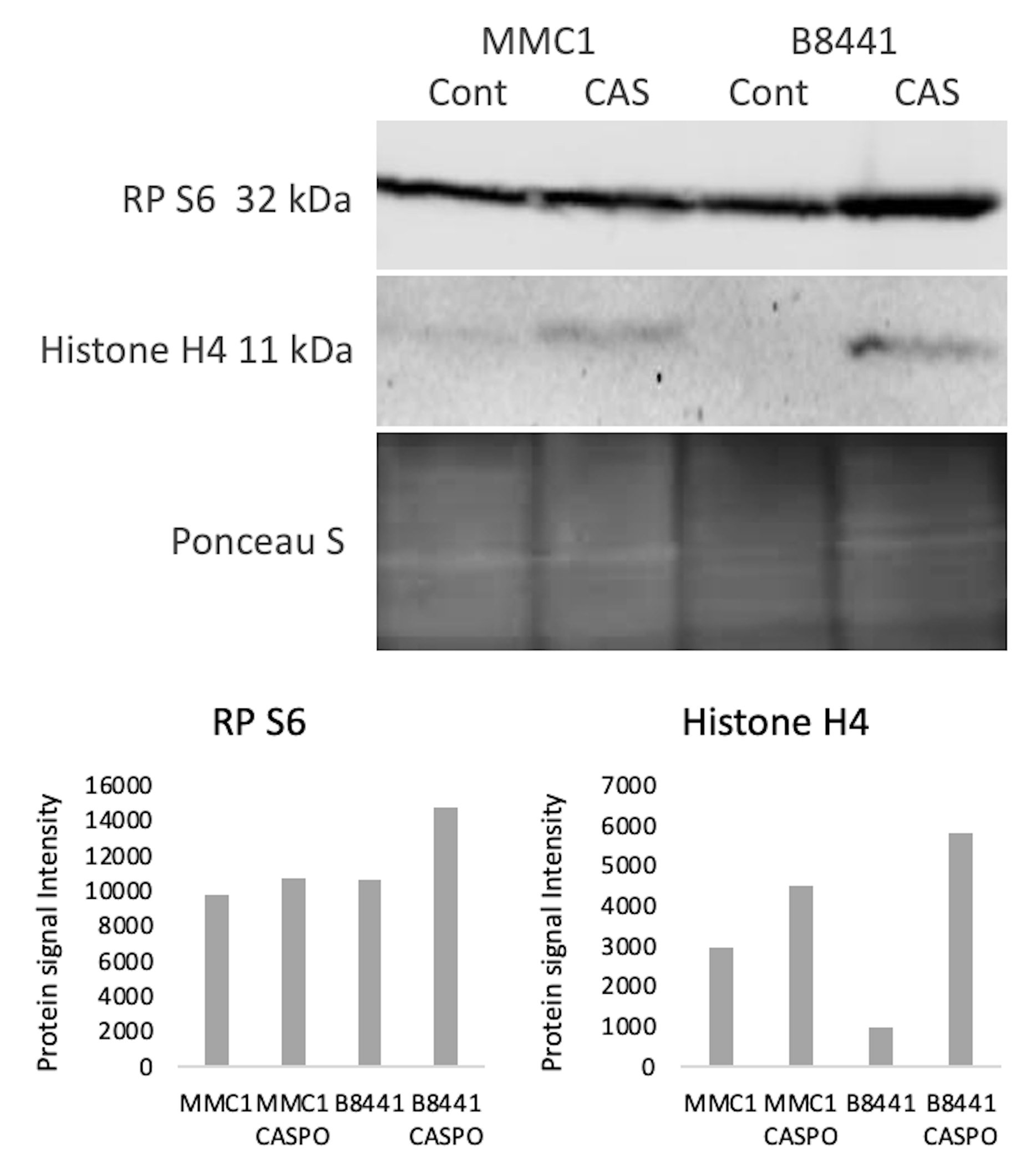

**Supplemental Figure 2**. Expression levels of Histone 4 and Ribosomal protein S6 for transcript level validation. Mouse anti-S6 700x dilution, rabbit anti-H4 800x dilution. For the secondary antibodies IRDye 680 10.000x dilution. The third lane refers to Ponceau staining to show the samples loading (30 ug of protein extract per lane). The bands were quantified by ImageJ, and the graphs represent the signal intensity (Y axis).

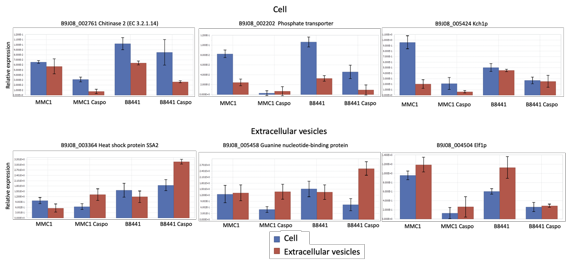

**Supplemental Figure 3**. Validation of the mRNA identified in *C. auris* yeast cells and EVs treated with caspofungin. Quantitative RT-PCR of mRNAs enriched in the Cell compared to the EVs (top). Transcripts more abundant in the EVs compared to the cell (bottom). Reverse transcription-qPCR was performed according to MIQE guidelines. The relative expression of transcript levels was normalized to the corresponding level of the histone H2A transcript and the error bars represent the mean and standard error of quadruplicate samples performed twice.
